# Sleep Disruption in a Mouse Model of Chronic Traumatic Brain Injury

**DOI:** 10.1101/2023.11.10.566553

**Authors:** Andrew R. Morris, Erwin K. Gudenschwager Basso, Miguel A. Gutierrez-Monreal, Rawad Daniel Arja, Firas H. Kobeissy, Christopher G. Janus, Kevin K. W. Wang, Jiepei Zhu, Andrew C. Liu

## Abstract

Chronic sleep/wake disturbances are strongly associated with traumatic brain injury (TBI) in patients and are being increasingly recognized. However, the underlying mechanisms are largely understudied and there is an urgent need for animal models of lifelong sleep/wake disturbances. The objective of this study was to develop a chronic TBI rodent model and investigate the lifelong chronic effect of TBI on sleep/wake behavior. We performed repetitive midline fluid percussion injury (rmFPI) in four months old mice and monitored their sleep/wake behavior using the non-invasive PiezoSleep system. The sleep/wake states were recorded before injury (baseline) and then monthly thereafter. We found that TBI mice displayed a significant decrease in sleep duration in both the light and dark phases, beginning at three months post-TBI and continuing throughout the study. Consistent with the sleep phenotype, these TBI mice showed circadian locomotor activity phenotypes and exhibited reduced anxiety-like behavior. TBI mice also gained less weight, and had less lean mass and total body water content, compared to sham controls. Furthermore, TBI mice showed extensive brain tissue loss and increased GFAP and IBA1 levels in the hypothalamus and the vicinity of the injury, indicative of chronic neuropathology. In summary, our study identified a critical time window of TBI pathology and associated circadian and sleep/wake phenotypes. Future studies should leverage this mouse model to investigate the molecular mechanisms underlying the chronic sleep/wake phenotypes following TBI early in life.

## INTRODUCTION

Sleep/wake disturbances (SWD) are among the most prevalent long-term consequences in patients suffering from traumatic brain injury (TBI).^1,2^ The Centers for Disease Control and Prevention (CDC) estimates that 1.6 to 3.8 million TBIs occur each year in the United States, many of which are left untreated.^3^ About 30-70% of patients following a mild TBI experience some form of SWD.^4,5^ Chronic post-TBI symptoms such as anxiety, headaches, irritability, and SWD persist in an estimated 43% of TBI patients that required hospitalization. Long-term sleep and circadian disruptions negatively affect life quality and have been associated with diseases, including cancer, diabetes, obesity, hypertension, daytime fatigue, anxiety, and depression, as well as cognitive deficits.^6–9^ SWD are known to contribute to morbidity and long-term health consequences and delay and impair TBI recovery; however, the underlying mechanisms are not well understood.^10–15^

While it is well recognized that TBI early in life is strongly associated with chronic SWD in patients, long-term preclinical models and studies are lacking.^16–18^ Lifelong SWD in mouse models has been understudied. The current study aimed to develop the first lifelong chronic TBI model with a central focus on SWD. Through the development of this model, future studies will offer mechanistic insights and treatment options, which ultimately will promote awareness and patient care in the clinic.

In this study, we utilized the murine repetitive midline fluid percussion injury (mFPI) model to investigate the long-term effects of TBI on the sleep/wake cycle and associated neuropathology.^19–21^ mFPI is one of the most widely used and extensively characterized preclinical TBI models.^22,23^ Our data brought light to the chronic TBI sequela influencing circadian rhythms and sleep/wake cycle and may offer new treatment strategies to improve TBI recovery.

## MATERIALS AND METHODS

### Animals

All experiments were conducted in accordance with institutional guidelines and approved by the University of Florida Institutional Animal Care and Use Committee (IACUC). Four months old male C57BL6 mice (Jackson Laboratories, Bar Harbor, ME) were used for the experiments. All mice were housed under controlled laboratory conditions with a 12-h light:dark schedule (lights on at 07:00 and lights off at 19:00), *ad libitum* access to food and water, temperature 20-26°C, and humidity 30-70%.

### Repetitive Midline Fluid Percussion Injury (rmFPI)

All rmFPI surgeries were performed as previously described.^24^ Mice were anesthetized with 4% isoflurane in 100% oxygen. Each mouse’s scalp was shaved and placed in a stereotactic frame fitted with a nose cone to maintain anesthesia with 2% isoflurane in 100% oxygen. A thermostatically controlled heating pad maintained the body temperature at 36.5°C to 37°C during the surgery. A midline sagittal incision exposed the skull from bregma to lambda. The skull was cleaned of periosteal connective tissue/fascia and dried using sterile cotton-tipped swabs and a 3.0 mm craniectomy was made midway between bregma and lambda along the sagittal suture without disrupting the underlying dura. The female portion of a Leur-Loc hub removed from a 20 gauge needle was affixed to the craniectomy site using cyanoacrylate. Dental acrylic was then applied around the hub to provide stability. After the dental acrylic hardened, the scalp was sutured around the hub, topical bacitracin ointment was applied to the incision site, and the animal was removed from anesthesia and monitored in a warmed cage until fully ambulatory. For the induction of injury, each animal was re-anesthetized with 4% isoflurane in 100% oxygen for two minutes and the male end of a spacing tube was inserted into the hub. The female end of the hub spacer assembly, filled with normal saline, was attached to the end of the fluid percussion apparatus (Custom Design and Fabrication, Sandston, VA). An injury of mild severity (1.62 ± 0.08 atmospheres) was administered by releasing a pendulum onto a fluid-filled cylinder to induce a brief fluid pressure pulse upon the intact dura. The pressure pulse measured by the transducer was displayed on a storage oscilloscope (Tektronix TDS2012C), and the peak pressure was recorded. After the injury, the animals were visually monitored for recovery of spontaneous respiration and a non-toxic, fast-drying silicone sealant (World Precision Instruments, Sarasota, FL) was applied to the hub to prevent contamination. Additional injuries were repeated at 24 hours and 48 hours after the initial injury respectively. Following recovery from the third injury, the hub and dental acrylic were removed and the incision was sutured under anesthesia. Topical bacitracin was then applied to the closed scalp incision. To alleviate postsurgical discomfort, sustained-release buprenorphine (Wedgewood Pharmacy, Swedesboro, NJ) was subcutaneously injected at the time of surgery and once every 48 hours. The duration of transient unconsciousness was determined by measuring the time it took each animal to recover the righting reflex as described before.^25^ After recovery of the righting reflex, animals were placed in a warmed holding cage to ensure the maintenance of normothermia and monitored during recovery before being returned to the vivarium. For animals receiving a sham injury, all of the above steps were followed with the exception of the release of the pendulum to induce the injury.

### Sleep Recordings

The non-invasive piezo sleep cage system (Signal Solutions, Lexington, KY) used in this study consisted of 16 individual units which simultaneously monitor sleep and wake states, as previously published.^26^ This system allows for longitudinal sleep/wake monitoring with greater throughput than EEG/EMG. PiezoSleep can assess total, day, and night sleep and wake durations, number of sleep bouts, and sleep bout length. Briefly, sleep was characterized by periodic (3 Hz) and regular amplitude signals recorded from the PVDF sensors, typical of respiration from a sleeping mouse. In contrast, signals characteristic of wake were both the absence of characteristic sleep signals and higher amplitude, irregular spiking associated with volitional movements. The piezoelectric signals in 2-second epochs were classified by a linear discriminant classifier algorithm based on frequency and amplitude to assign a binary label of ‘sleep’ or ‘wake’. Mice sleep in a polycyclic manner (more than 40 sleep episodes per hour) and so mouse sleep was quantified as the minutes spent sleeping per hour, presented as a percentage for each hour. Data collected from the cage system were clustered into bins over specified periods (e.g. 1 hour) using the average of percentage sleep, as well as binned by the length of individual bouts of sleep, and the mean bout lengths were calculated. Sleep data were collected monthly. Hourly percentage sleep was calculated by averaging the percentage of sleep for all days of a given collection by a blinded investigator.

### Body Composition

Body composition was assessed by nuclear magnetic resonance in conscious mice using the EchoMRI whole body composition analyzer (EchoMedical Systems), as previously described.^27^

### Elevated Plus Maze (EPM) Test

The EPM apparatus consisted of two darkened and walled (but uncovered) arms and two exposed arms joined by a central square (75.5 cm × 75.5 cm × 6 cm), elevated 50.5 cm above the floor. Mice were individually placed in the center square (facing the north closed arm) and activity was tracked for five minutes. A camera placed above the maze recorded the animal’s movement and was analyzed using ANY-maze 7.20 (Stoelting).

### Wheel-Running Locomotor Activity Assay

Mice were individually housed in cages equipped with running wheels and locomotor activity was recorded as we’ve previously described.^28,29^ The mice were acclimated to the running wheel cages under a standard 12hr/12hr light/dark cycle for two weeks and then released to constant darkness (DD) for two weeks. The wheel-running activity was recorded using the ClockLab 4.011 program (Actimetrics) and analyzed using the ClockLab Analysis 6.0.52 program (Actimetrics).

### Histological Processing, Staining, and Lesion Volume

Mice were transcardially perfused using a mini-peristaltic pump (Fisher Scientific, Pittsburgh, PA) with ice-cold phosphate-buffered saline (PBS) followed by 4% paraformaldehyde (PFA) as previously described.^30^ Perfused brains were post-fixed in 4% PFA and then cryoprotected overnight in 15% sucrose followed by 30% sucrose overnight at 4°C. Brains were embedded in a solution of 30% sucrose in Optimal Cutting Temperature (OCT) medium and stored at -80°C. Tissue sections of 30 µM thickness and 600 µM apart were sectioned using a Leica cryostat (Leica Biosystems, Wetzlar, Germany), starting at -0.5 anterior-posterior (AP) bregma level up to 2.5 AP bregma level.

For the detection of Nissl substance in the cytoplasm of neurons, sections were stained with 0.1% cresyl violet acetate (Electron Microscopy Sciences, Hatfield, PA) for 30 minutes, following standard methodologies. Scanned coronal sections were analyzed using FIJI v1.54f to measure the percentage of brain tissue loss. Total healthy brain area in (*HB*) and injury area (*IA*) was calculated to determine the percentage of tissue lost by the impact:

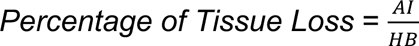

Healthy brain was characterized by normal architecture and neuronal density after cresyl violet staining. In contrast, injured tissue was determined by loss of neuronal density, and tissue reabsorption after chronic TBI. Borders between healthy and injured tissue were normally sharp and clearly defined. Injury as a percentage of brain area was calculated by adding the uninjured area. Lesion volume (*LV*) was calculated as the volume (mm^3^) of a sphere with the estimated injured area:

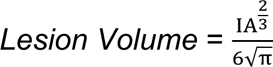

For immunofluorescence detection, sections were washed in PBS then placed in boiling sodium citrate buffer (10mM sodium citrate, 0.05% Tween 20, pH 6.0) for 20 minutes for antigen retrieval. Tissue was washed with PBS and blocked (5% serum, 0.4% Triton X-100 in PBS) for one hour at room temperature. Sections were then incubated in a humid chamber overnight at 4°C in primary antibody: anti-GFAP (Abcam [ab4674], 1:200 dilution), anti-Iba1 (Abcam [ab178846], 1:1000 dilution). Next, tissue was washed in PBS and incubated in fluorochrome-conjugated secondary antibody for one hour at room temperature (Abcam [ab150076, ab150173], 1:500 dilution). Sections were then washed in PBS and mounted and cover-slipped in aqueous mounting media containing DAPI.

For Nissl body staining, GFAP, and Iba1, slides were imaged using a Keyence BZ-X800 microscope (Keyence Corporation, Osaka, Japan) at 4X magnification. GFAP and Iba1 stained slides were also imaged using a ZEISS inverted confocal microscope (ZEISS, Oberkochen, Germany) at 10X or 20X magnification. Images were processed using ZEN lite 3.8 (ZEISS, Oberkochen, Germany).

### Western Blot

Mice were euthanized by isoflurane overdose followed by cervical dislocation and brains were dissected in PBS over ice. The sagittal cortex surrounding the injury site and hippocampus were rapidly collected, frozen in liquid nitrogen, and stored at -80°C. Tissues were homogenized with micropestle and motor in 1X diluted RIPA buffer containing protease inhibitors. Lysate was centrifuged and the supernatant was collected for protein quantification. Protein extracts were resolved on a 12% SDS-PAGE gel and transferred to a nitrocellulose membrane using the iBlot® (Invitrogen) gel transfer device. Membranes were incubated with antibodies against GFAP (cat # RPCA-GFAP, EnCor Biotechnology) and alpha fodrin/alpha-II spectrin (cat # BML-FG6090-0500, Enzo Life Sciences). Protein bands were visualized using horseradish peroxidase-conjugated anti-mouse or anti-rabbit IgG (Abcam) and ECL (Thermo Fisher Scientific). Beta-actin detection was used as an internal quantitative control, expression levels of targeted proteins were normalized those of each investigated protein in the densitometry analysis. Positive signals were detected using the ChemiDoc (Bio-Rad Laboratories) imaging system. Densitometric analyses were performed using ImageJ (National Institutes of Health).

### Statistical Analysis/Evaluation

Data are shown as the mean ± standard error of the mean (SEM) and analyzed using statistical software (GraphPad Prism 9). A *p*-value < 0.05 was considered statistically significant. A two tailed t-test was used to compare the sham and TBI groups.

## RESULTS

### TBI induced a transient loss of righting reflex and reduction in weight gain

Following pre-injury baseline sleep recording, 20 age-matched mice were randomly assigned to either sham or TBI groups (Fig. 1). Mice were acclimated until four months of age, which is considered the mature adult life phase equivalency.^31^ Surgery was performed in the morning and mice were allowed to recover for two to four hours before being subjected to three mFPIs or sham injuries in the afternoon. Each mouse was injured every 24 hours apart following the previous TBI or sham injury. There were no differences in FPI pressure between the first, second, and third injuries (1.60 ± 0.04 atm; 1.60 ± 0.11 atm; 1.65 ± 0.08 atm). Following the final injury, mice were allowed to recover for two weeks before beginning monthly sleep recordings in a longitudinal sleep study (Fig. 1).

**FIG. 1.**
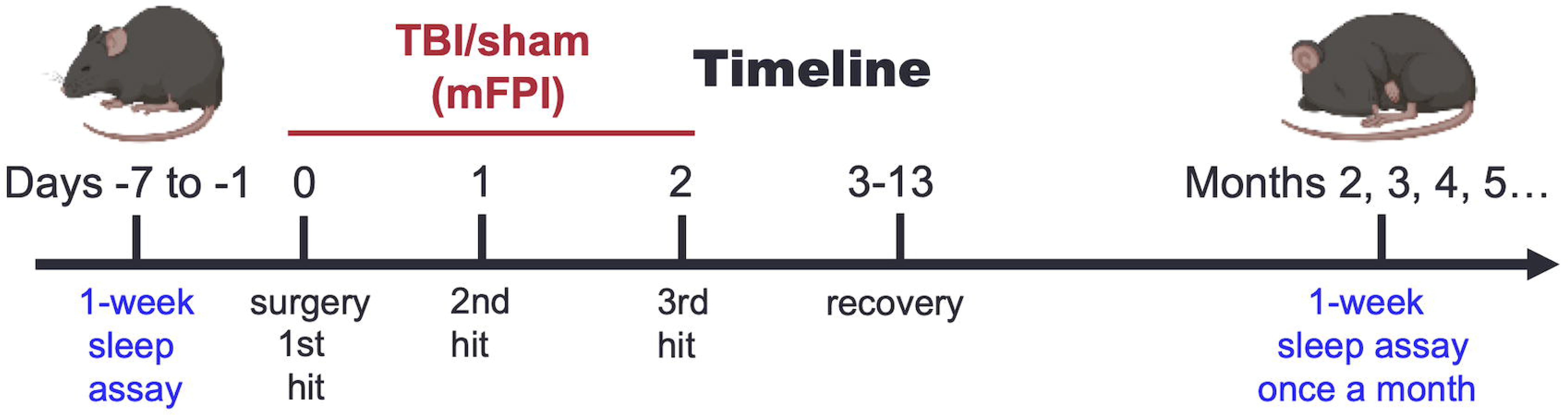
Timeline for chronic sleep evaluation in the repetitive midline fluid percussion injury (rmFPI) model. Mice were randomly assigned to either the TBI or sham groups. Baseline sleep was recorded for all mice prior to surgery. rmFPI or sham injury was performed three times at 24-hour intervals following craniectomy and placement of the Luer hub. Following recovery, sleep was recorded monthly.

The righting reflex is the innate tendency of the mouse to return upright after being placed on its side or back. The time duration for loss of righting reflex (LRR) after injury/sham and discontinuation of anesthesia is comparable to the duration of lost consciousness in TBI patients.^32^ LRR was used as an index of injury severity in this study as used before.^33–36^ On the first day of injury, LRR was considerably longer (366 ± 53 seconds) in rmFPI mice than in shams (138 ± 110) (Fig. 2A). Similarly, on the second day of injury, TBI mice LRR was 166 ± 93 seconds longer than shams (rmFPI 314 ± 93 seconds; sham 148 ± 92 seconds) (Fig. 2A). On the third and final day of injury, TBI LRR was 244 ± 97 seconds while sham mice LRR was 173 ± 69 seconds (Fig. 2A). Overall, sham mice demonstrated a shorter LRR compared to rmFPI mice despite the injury being the same, indicative of the immediate effect of brain trauma and confirming injury in the TBI group as reported in previous studies.^33–36^

**FIG. 2.**
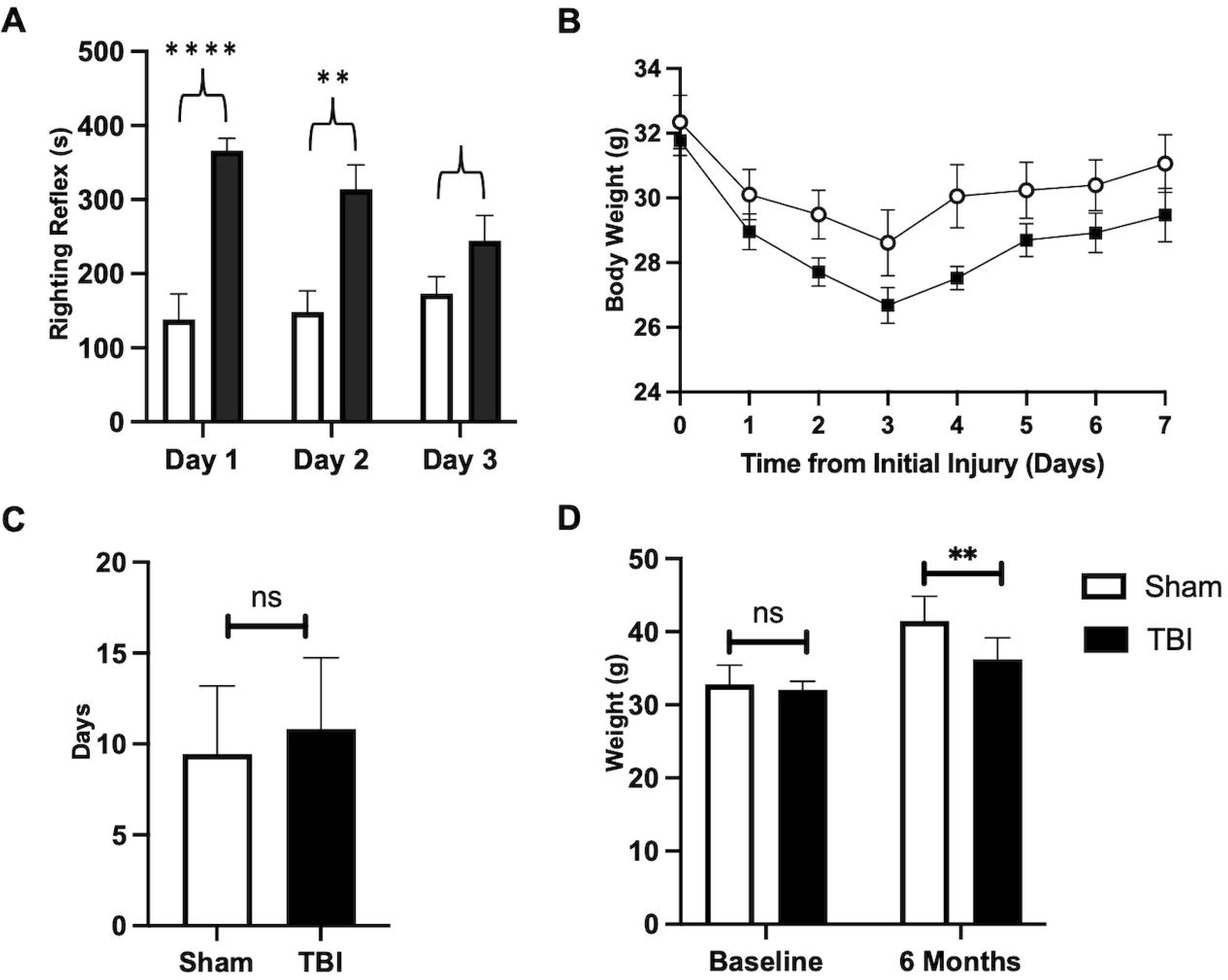
Repetitive midline fluid percussion injury (rmFPI) induced changes in righting reflex and body weight. (**A**) rmFPI induced a transient loss of righting reflex following each injury. (**B-C**) Daily body weight during the first week after injury. TBI mice recovered to baseline body weight more slowly than sham controls. (**D**) rmFPI mice gained less weight compared to shams six months after injury. * *p* < 0.05; ** *p* < 0.01; **** *p* < 0.0001; ns, not significant.

To evaluate recovery after trauma and overall health condition, we recorded body weight at baseline and daily following injury until mice recovered back to baseline weight (Fig. 2B). On average, rmFPI mice required more days to recover to baseline body weight compared to sham controls (Fig. 2C). Evaluation of body weight six months post-injury revealed that rmFPI mice gained less weight relative to shams despite sharing the same baseline body weight, indicating the long-term effect of TBI on systemic metabolism (Fig. 2D).

### TBI mice exhibited sleep deficits beginning at three months post-injury

To evaluate the chronic effect of rmTBI on sleep/wake behavior, we used the non-invasive piezoelectric sleep cage system for longitudinal sleep monitoring.^16^ Beginning at three months post-injury, a phenotype in sleep duration was observed (Fig. 3A, 3B, 3C). More specifically, significant hourly differences were observed at zeitgeber time (ZT) 10 (during the light phase) as well as ZT13, ZT14, ZT15, and ZT20 (during the dark phase) where rmTBI mice slept less than sham controls (Fig. 3A). Sleep duration was decreased in rmTBI mice during the light phase, dark phase, and total compared to sham controls (Fig. 3B, 3C). At three months post-injury, rmTBI mice spent 430 minutes sleeping during the light phase compared to 467 minutes by sham controls (Fig. 3B, left panel). During the dark phase, rmTBI mice slept 201 minutes compared to 255 minutes by sham controls (Fig. 3B, right panel). In total, rmTBI mice spent 631 minutes sleeping compared to 722 minutes by sham controls; a difference of 91 minutes per day (Fig. 3C). Prior to three months post-injury, there were no significant differences between groups in the light phase, dark phase, or total sleep amounts (Fig. 3C). Total sleep differences between sham controls and rmTBI mice remained significant on a monthly basis for the remainder of life post-TBI (Fig. 3C).

**FIG. 3.**
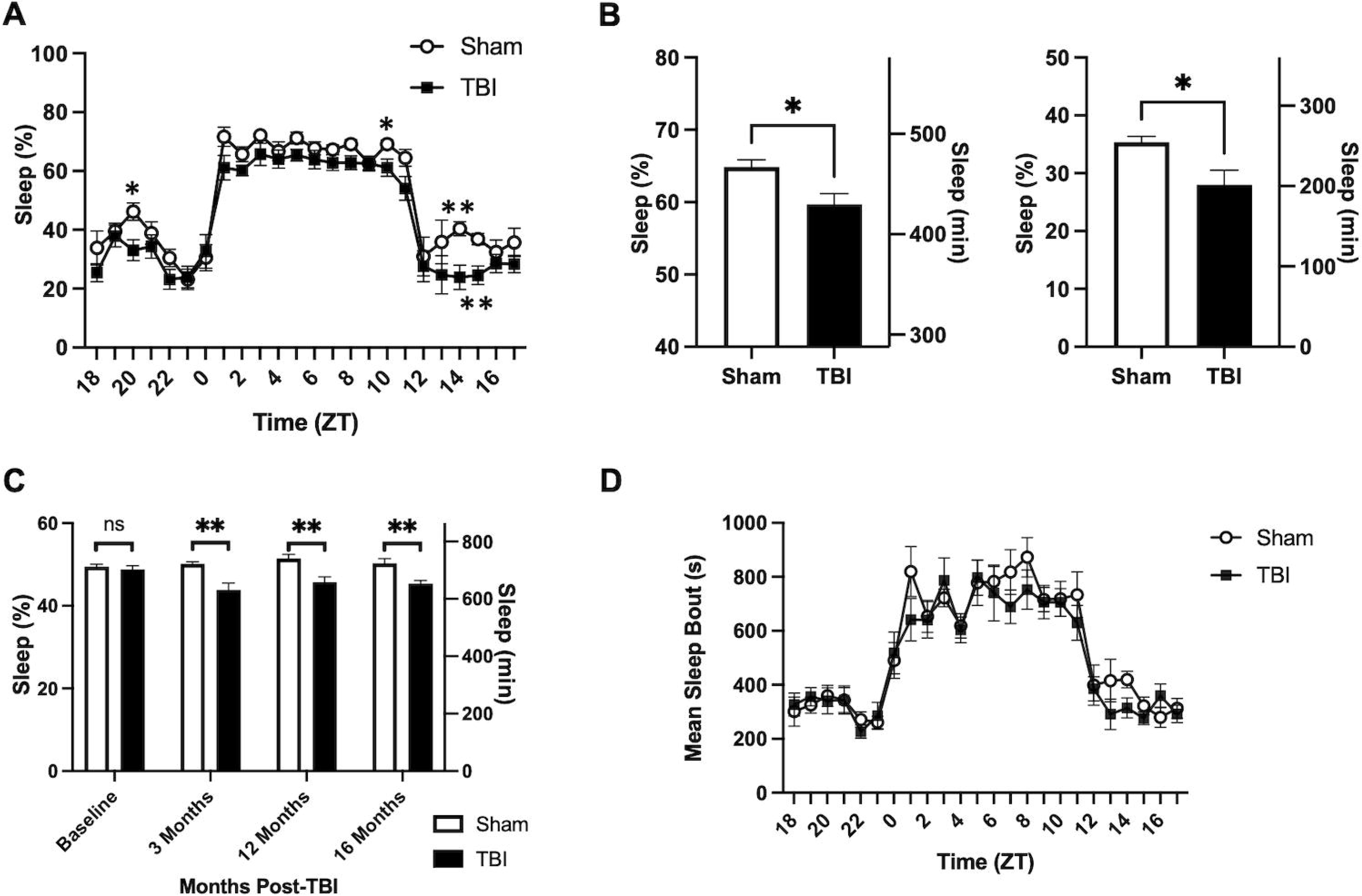
Repetitive midline fluid percussion injury (rmFPI) mice exhibited sleep disturbances starting at three months post-injury. **(A)** Hourly sleep percentage at three months post-injury. Sleep was recorded in a non-invasive piezoelectric sleep system. ZT, Zeitgeber time. rmFPI mice showed sleep reduction at ZT10, ZT13, ZT14, ZT15, and ZT20, compared to shams. (**B)** Sleep percentage and amount at three months post-injury. rmFPI mice slept less than sham controls in both the light and dark phases. **(C)** Monthly sleep duration. Compared to sham controls, rmFPI mice began to show sleep reduction at three months and the phenotype continued through 16 months post-TBI. **(D)** Hourly mean sleep bout lengths at three months post-injury. * *p* < 0.05; ** *p* < 0.01; ns, not significant.

At three months post-injury, mean sleep bouts were near significance (*p*=0.0569) at ZT14 with rmTBI mice averaging 315 ± 116 second sleep bouts compared to 420 ± 82 second sleep bouts for sham controls (Fig. 3D). Mean sleep bouts were lower during the light phase (rmFPI 684 ± 129 seconds; sham 727 ± 71 seconds), dark phase (rmFPI 316 ± 58 seconds; sham 334 ± 46 seconds), and total (rmFPI 500 ± 75 seconds; sham 531 ± 33 seconds) for rmTBI compared to sham controls, but not statistically significant. By 16 months post-injury, sleep quality during the light phase worsened (rmFPI 676 ± 207 seconds; sham 720 ± 170 seconds) but did not reach statistical significance.

### TBI had a long-term effect on circadian locomotor activity

The sleep/wake cycle is regulated by the central circadian clock located in the suprachiasmatic nucleus (SCN) of the hypothalamus. As a clock output measurement, circadian wheel-running activity generally reflects the function of the SCN clock.^37,38^ In the presence of a light-dark cycle, mice are generally most active during the dark phase and relatively inactive during the light phase. The endogenous circadian rhythms persist under free-running conditions such as constant darkness and can be used to determine the circadian behavioral phenotypes. At 17 months post-injury, wheel-running behavior was assessed in mice under constant darkness (Fig. 4A). TBI mice demonstrated a significant increase in wheel-running activity at circadian time (CT) 12 and at CT18-20. This result suggests that TBI had a long-term effect on the SCN clock and influenced circadian locomotor behavior.

**FIG. 4.**
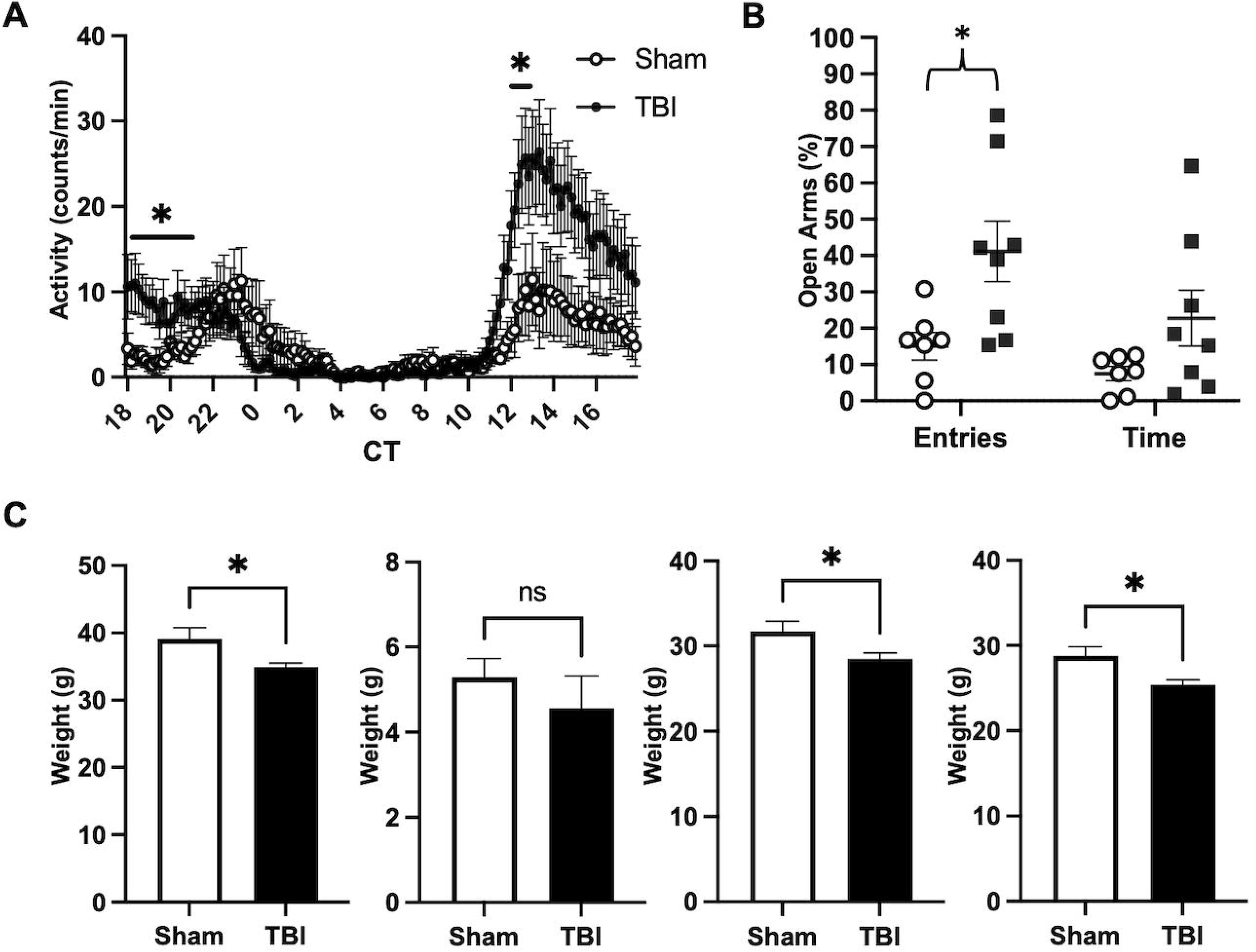
Repetitive midline fluid percussion injury (rmFPI) led to changes in behavior and body composition at 17 months post-injury. (**A**) Wheel-running activity in rmFPI mice at 17 months post-injury was significantly increased at CT 12-13 and CT18-21 relative to shams. (**B**) Behavioral tests showed that rmFPI mice had an increase in the percentage of entries into the open arms at the elevated plus maze test. (**C**) Body weight and composition. rmFPI mice weighted less than their controls at 17 months post-injury, similar to at 6 months (see Figure 2). EchoMRI body composition analysis showed no significant changes in fat amount, but a reduction in lean mass and total water in rmFPI mice relative to shams. * *p* < 0.05; ns, not significant.

### TBI mice displayed behavioral disinhibition

Anxiety and depression are psychiatric sequelae often associated with SWD in clinical TBI patients.^39^ Here, we used the Elevated Plus Maze to examine anxiety-like behavior 13 months after the sleep phenotype began. TBI mice entered anxiogenic zones with greater frequency compared to sham controls. The risk-taking behavior is likely due to increased impulsivity. At 16 months post-injury, anxiety-like behavior was measured in mice using the elevated plus maze. rmTBI mice made a significantly higher proportion of entries into the open arms of the apparatus (41.1 ± 23.7%) compared to sham controls (15.0 ± 9.9%) (Fig. 4B). The proportion of time spent in anxiogenic zones was marginally increased in TBI mice (32.9 ± 28.4%) compared to sham controls (12.4 ± 8.9%). While rmTBI didn’t exhibit increased anxiety-like behavior that was observed in other TBI models, they did demonstrate greater risk-taking behavior indicative of a behavioral disinhibition phenotype.^40^

### TBI led to changes in body composition later in life post-injury

Body weight discrepancy between rmFPI mice and sham controls persisted at 17 months post-injury (Fig. 4C, left panel). Considering the body mass reduction in rmFPI mice, we measured mice body composition using the EchoMRI system (Fig. 4C). There were no significant differences in fat mass content between the groups (Fig. 4C, left center panel). However, rmFPI mice showed a decrease in lean mass compared to sham controls (Fig. 4C, right center panel). Lean mass is a muscle tissue mass equivalent to all the body parts containing water, excluding fat and bone minerals. This reduction in lean mass may indicate atrophy of muscle or other organs.^41^ In addition, rmFPI mice exhibited a lower total water content compared to sham controls (Fig. 4C, right panel). These results suggest that the weight reduction in rmFPI mice does not involve a reduction in fat storage but is likely attributed to muscular and organ atrophy.^42,43^

### TBI induced chronic lesioning and neuropathology

To evaluate the long-term effect of rmFPI on brain morphology, we performed perfusion and histological analysis of brain sections. Gross inspection of rmFPI brains 19 months post-injury revealed extensive cavitation (Fig. 5A). Affected areas included both hemispheres of the parietal-temporal lobes and part of the frontal lobes, corresponding to the retrosplenial cortex, primary motor cortex, and primary somatosensory cortex. In contrast, sham brains showed no signs of trauma. Cresyl violet coronal sections showed an extension of the injury penetrating up to the thalamus and the anterior areas of both hippocampus (Fig. 5B, left panel). Compared to sham controls, a significant percentage (17.75%) of the rmFPI brain tissue was lost due to injury (Fig. 5B, middle panel). Volumetric estimation showed that the injury was 31.48 mm^3^ for rmFPI brains and only 0.146mm^3^ for sham (Fig. 5B, right panel). These data showed clear differences of histopathology between rmFPI and sham brains. Previous mFPI studies in mice at early time points showed no clear brain tissue loss at gross examination or histological evaluation.^20,44^ To our knowledge, this is the first study that extends to more than one year after injury.

**FIG. 5.**
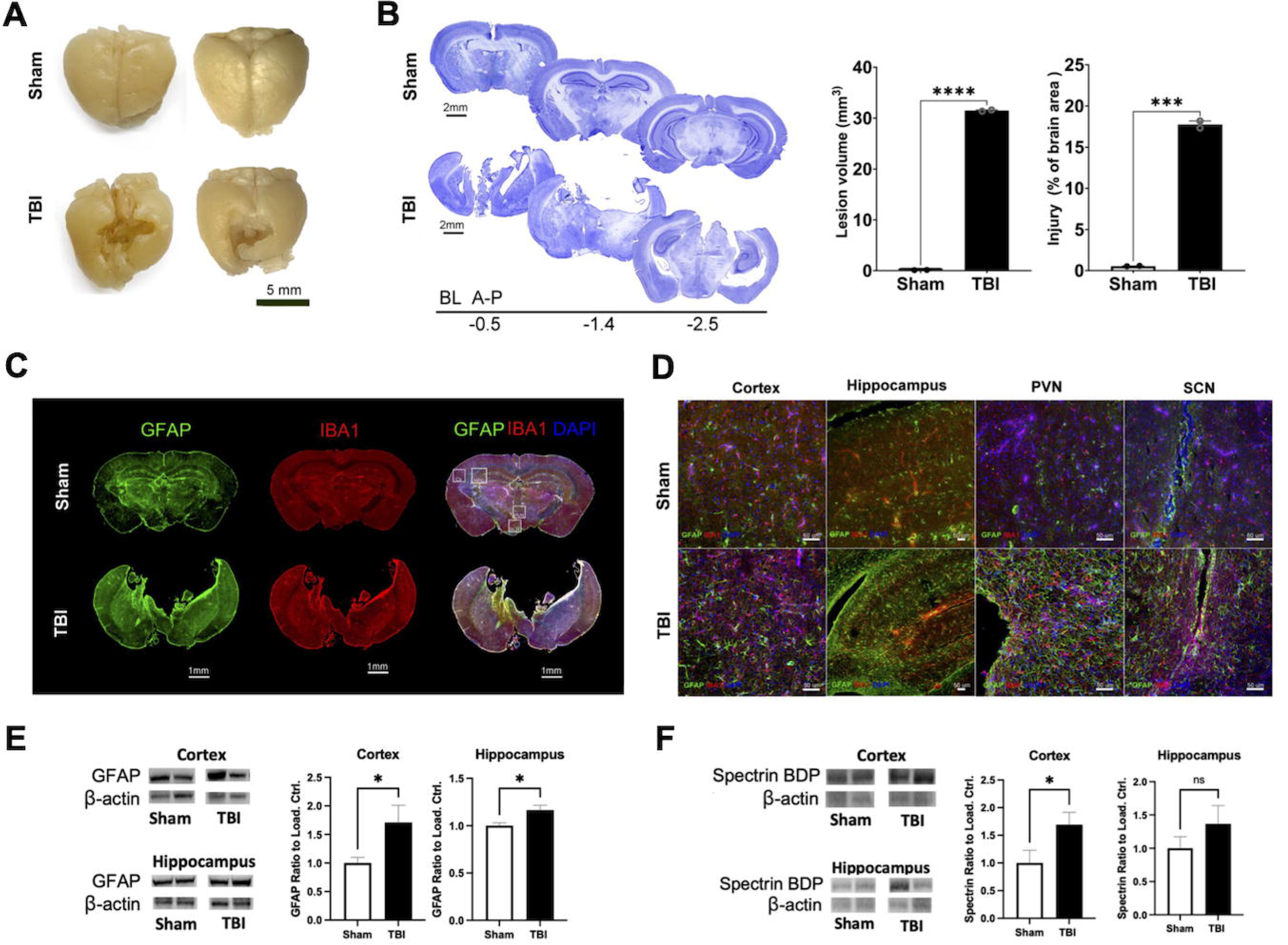
Repetitive midline fluid percussion injury (rmFPI) induced damage of cortex and hippocampus associated with glial activation. (**A**) Gross image of post-mortem brains showing extensive cavitation. (**B**) Cresyl violet coronal brain sections showing that the injury extended to the thalamus and hippocampus. Quantification of the lesion (31.48 mm^3^) represents 17.75% of the brain area of rmFPI mice, compared to 0.146mm^3^ for sham controls. (**C**) Immunofluorescence images of GFAP and IBA1 staining. rmFPI brains showed an increase in GFAP and IBA1 levels, especially in areas adjacent to the injury site. (**D**) Confocal images revealed an increase in GFAP and IBA1 signals in the cortex, hippocampus, PVN, and SCN of rmFPI brains compared to shams. PVN, paraventricular nucleus; SCN, suprachiasmatic nucleus. (**E-F**) Western blot analysis of GFAP (**E**) and alpha-II spectrin (**F**) in the cortex and hippocampus of sham and TBI mice. BDP, breakdown product; * *p* < 0.05; *** *p* < 0.001; **** *p* < 0.0001; ns, not significant.

To determine the degree of TBI-induced histopathological alterations in the brain, we used stained tissue sections against IBA1 and GFAP to evaluate microglia/macrophage and astrocyte changes.^45,46^ Microglial activation has been linked to sleep dysregulation in mice and humans and GFAP plays a critical role in astrogliosis following injury and neurodegeneration.^47,48^ We observed an increase in GFAP and IBA1 levels in rmFPI brains, especially in areas adjacent to tissue loss (Fig. 5C). Higher magnification confocal images revealed an increase in GFAP and IBA1 signals in the cortex, hippocampus, and hypothalamus (more specifically, the PVN and SCN, known to regulate circadian rhythms and the sleep/wake cycle) of rmFPI brains (Fig. 5D).^49–52^ Thus, the long-term effect of TBI on the hypothalamus may explain the sleep changes in these rmFPI mice. Western blot analysis of GFAP level in the cortex and hippocampus further confirmed this finding (Fig. 5E). Furthermore, our Western blot data also showed that alpha-II spectrin breakdown product (BDP) was significantly increased in the cortex and slightly increased in the hippocampus of TBI mice (Fig. 5F). Alpha-II spectrin is abundant in neurons and cleavage into 145/150 kDa BDP by calpain/caspase is indicative of TBI-induced cell death.^53,54^ Overall, these results show that several brain regions are altered 18 months post-injury and the pathology data support the sleep phenotypes seen in TBI mice.

## DISCUSSION

TBI is a serious public health problem that impacts athletes, civilians, and military personnel worldwide. With reported cases on the rise in recent years, TBI is an important health concern that impacts patients’ quality of life over their lifespan.^55^ Mild TBIs are the most common, representing approximately 80% of cases.^56^ SWD are one of the most prevalent chronic sequelae and, despite occurring in upwards of 70% of mild TBI cases, are underrecognized and not well studied.^57^

While many murine studies have investigated sleep following TBI, the focus has been on the acute phase and not the chronic timepoints.^19,58^ To our knowledge, this is the first longitudinal study of chronic SWD in mice. Interestingly, a previous sleep study using mFPI concluded that a diffuse axonal injury model does not lead to chronic SWD in mice.^19^ However, that study only studied sleep behavior for five weeks with a single mFPI and suggested that a secondary injury may be necessary for the induction of chronic sleep disruption in mice. Sleep behavior phenotypes did not develop in our model of rmFPI until three months post-injury. This decreased sleep quantity phenotype persisted throughout the lifespan of the TBI mice. Circadian locomotor activity pattern was consistent with sleep/wake behavior. TBI mice slept less than sham controls and were increasingly active.

It has been hypothesized that sleep is regulated by the central circadian clock located in the SCN and the homeostatic process (whose precise anatomical location is not known).^59,60^ While the hypothalamus is one of the key sleep homeostatic centers, the SCN is also known to be involved in the regulation of sleep homeostasis. The circadian clock also regulates immune function and recent studies have established a bidirectional relationship.^50,61,62^ Further, recent studies from our group have shown that the proinflammatory NF-kB transcription factor directly interacts with the core clock component BMAL1 and interferes with the circadian transcriptional feedback mechanism.^62^ Following TBI, NF-kB is activated within hours in neurons, but it is not detectable after two weeks.^63^ Similarly, constitutively active NF-kB in neurons leads to circadian disruption and sleep dysfunction.^62,64^ Therefore, it is likely that neuronal NF-kB expression could play a role in the acute TBI effects, but its role in the chronic effects is not clear. Meanwhile, NF-kB activation in microglia has been shown to appear within 24 hours after injury but persists long-term.^63,65,66^ Microglia, as the primary innate immune response, become “primed” following TBI incident and elicit a hyperreactive response to subsequent challenges.^67,68^ Previous studies have suggested that a secondary challenge may be necessary for chronic sleep and behavioral dysfunction.^19,68^ These long-term neuropathological interactions that disturb sleep through the SCN clock and the hypothalamic homeostatic center are not well understood and warrant future neurobehavioral and mechanistic studies.

Previous studies using repetitive models of TBI have demonstrated a similar trend of shortened loss of righting reflex times after successive injuries.^69^ In our study, no significant body weight differences between groups were observed at baseline. However, body weight decreased after each successive injury for both sham and TBI mice but began to recover 48 hours following the third and final hit. While TBI mice (on average) required an additional day to recover to baseline weight, body weight remained significantly below that of sham controls over their lifespan. Fat, lean, free water, and total water body composition was measured to further explore these differences. While group housing is the preferred default in the laboratory setting, single housing occurs when necessary to prevent prolonged aggression and severe injury to mice, especially in males.^70^ However, single housing has been shown to decrease body mass and increase adiposity, compared to group housing.^71–74^ Therefore, single-housed mice were removed from the body composition assessment. Body composition was measured at the end of life due to chronic differences noted in body weight. While there were significant differences in fat and free water, TBI mice had less lean mass and total water compared to sham controls.

A limitation of this study was the usage of only male mice and did not explore sex-related differences in females. Future studies will also look at the chronic effects of rmFPI in females. Due to the longitudinal nature of the study, mechanistic approaches could not be utilized, and interventions were kept to a minimum through the end of life.

Furthermore, this study laid a foundation for future investigations into the underlying mechanisms and cause-and-effect relationship between TBI pathology and chronic SWD. Having identified specific time points at which SWD emerge, further studies can focus on these critical time windows to explore the molecular and cellular changes that contribute to these disruptions. Additionally, exploring new targets and therapeutic approaches to mitigate the SWD associated with TBI can enhance the overall well-being and quality of life for TBI patients.

Future studies should expand upon these long-term effects, exploring the impact of TBI on sleep in larger cohorts and potentially including diverse populations, including female mice and different TBI severity models. By delving deeper into the complex interactions between TBI pathology, sleep disruption, and associated behavioral and molecular changes, we can gain a more comprehensive understanding of the underlying mechanisms and develop targeted interventions for SWD in TBI patients.

In summary, this study highlights the critical time window of TBI pathogenesis associated with chronic SWD and provides valuable insights for future research. By shedding light on the relationship between TBI and SWD, this work paves the way for further investigations that will ultimately lead to improved long-term solutions for individuals living with TBI-related sleep disorders.

## CONCLUSIONS

In conclusion, this study identified a critical time window of TBI pathology, starting at three months post-injury, during which chronic SWD became apparent. By monitoring sleep/wake states in a chronic murine model of TBI, the study revealed a significant decrease in sleep duration during both the light and dark phases, persisting throughout their lifespan and accompanied by a deterioration in sleep quality. These findings underscore the importance of considering the long-term consequences of TBI on sleep. Future studies will focus on the most appropriate time window of pathophysiology, including the sleep phenotype onset. Future mechanistic studies could offer new targets for intervention, targeting SWD in TBI patients. TBI mice also demonstrated significant decreases in body weight and body composition compared to their sham injury controls. Future studies will explore feeding behaviors and the functional relevance of such differences.

## TRANSPARENCY, RIGOR AND REPRODUCIBILITY SUMMARY

The sample size per group was planned based on availability of non-invasive PiezoSleep cages, a limiting factor. A total of 20 mice survived the experimental manipulations. Mice were assigned to rmFPI or sham groups by random alternation. Analyses of experimental materials were performed by investigators blinded to relevant characteristics of the subjects. Replication by the study group is ongoing. All equipment and analytical reagents used to perform experimental manipulations and measurements are widely available.

## Acknowledgments

We would like to acknowledge funding from the National Institutes of Health (1R21AA029785-01A1). We would also like to express our gratitude to the University of Florida Claude D. Pepper Older Americans Independence Center (OAIC) Circadian Rhythms research core for allowing us to use the sleep and activity recording equipment. We would also like to thank Dr. Karyn A. Esser for use of the EchoMRI whole body composition analyzer and Drs. Belinda S. Pinto and Eric T. Wang for use of the ZEISS inverted confocal microscope.

## Author Contributions Statement

**Andrew Morris:** Conceptualization, Methodology, Formal analysis, Investigation, Data Curation, Writing – Original Draft, Writing – Review & Editing, Visualization, Project administration **Erwin K. Basso:** Methodology, Formal analysis, Data Curation, Writing – Review & Editing **Miguel A. Gutierrez-Monreal:** Writing – Review & Editing **R. Daniel Arja:** Investigation, Writing – Review & Editing **Firas H. Kobeissy:** Formal analysis, Writing – Review & Editing, Supervision **Christopher G. Janus:** Formal analysis, Resources, Writing – Review & Editing, Supervision **Kevin K.W. Wang:** Resources **Jiepei Zhu:** Resources, Investigation, Writing – Review & Editing **Andrew C. Liu:** Resources, Writing – Review & Editing, Supervision, Funding acquisition

